# Intrinsic activity temporal structure reactivity to behavioural state change is correlated with depressive symptoms

**DOI:** 10.1101/703496

**Authors:** Niall W. Duncan, Tzu-Yu Hsu, Paul Z. Cheng, Hsin-Yi Wang, Hsin-Chien Lee, Timothy J. Lane

**Affiliations:** Graduate Institute of Mind, Brain and Consciousness, Taipei Medical University, Taipei, Taiwan; Brain and Consciousness Research Centre, TMU Shuang-Ho Hospital, New Taipei City, Taiwan; Department of Psychiatry, School of Medicine, College of Medicine, Taipei Medical University, Taipei, Taiwan; Department of Psychiatry, TMU Shuang-Ho Hospital, New Taipei City, Taiwan

**Keywords:** Negative thought, self-focus, long-range temporal correlations, criticality

## Abstract

The brain’s intrinsic activity plays a fundamental role in its function. In normal conditions this activity is responsive to behavioural context, changing as an individual switches between directed tasks and task-free conditions. A key feature of such changes is the movement of the brain between corresponding critical and sub-critical states, with these dynamics supporting efficient cognitive processing. Breakdowns in processing efficiency can occur, however, in brain disorders such as depression. It was therefore hypothesised that depressive symptoms would be related to reduced intrinsic activity responsiveness to changes in behavioural state. This was tested in a mixed group of major depressive disorder patients (n = 26) and healthy participants (n = 37) by measuring intrinsic EEG activity temporal structure, quantified with detrended fluctuation analysis (DFA), in eyes-closed and eyes-open task-free states and contrasting between the conditions. The degree to which DFA values changed between the states was found to be negatively correlated with depressive symptoms. DFA values did not differ between states at all in those with high symptom levels, meaning that the brain remained in a less flexible sub-critical condition. This sub-critical condition in the eyes-closed state was further found to correlate with levels of maladaptive rumination. This may reflect a general cognitive inflexibility resulting from a lack in neural activity reactivity that may predispose people to overly engage in self-directed attention. These results provide an initial link between intrinsic activity reactivity and psychological features found in psychiatric disorders.

## Introduction

Ongoing brain activity appears to function close to an unstable equilibrium point (Deco and Jirsa, 2012). This critical state, where the system is on the boundary between order and disorder (Beggs and Timme, 2012), has been linked to efficient information processing and optimal behavioural performance (Irrmischer et al., 2018b; Palva et al., 2013; Palva and Palva, 2018). A system in a critical state is characterised by scale-free activity distributions and long-range temporal correlations (LRTC; Plenz and Thiagarajan, 2007), which can be measured with techniques such as detrended fluctuation analysis (DFA; Peng et al., 1995). Such approaches have been used to study brain activity in a variety of species and experimental conditions (Bellay et al., 2015; Priesemann et al., 2009; Ribeiro et al., 2010; Shriki et al., 2013).

This prior work suggests that the brain displays variability around a critical state depending upon the current behavioural condition (Bellay et al., 2015; Hahn et al., 2017). One such behavioural change is the switch from having closed eyes to having them open (Yan et al., 2009). This eyes-open (EO) effect is known to influence the activity in multiple brain networks, impacting upon neural oscillations within different frequency bands (Barry et al., 2007; Barry and De Blasio, 2017). Activity in the eyes-closed (EC) condition appears to be closer to criticality, moving towards sub-critical dynamics in the eyes-open condition (Hahn et al., 2017; Jao et al., 2012).

The ability to effectively shift between dynamic cortical states depending on behavioural context has been linked to cognitive performance (Irrmischer et al., 2018b). This may reflect an ability to move easily from a critical state, where there is the maximum potential to respond flexibly to changing demands, to a sub-critical one where processing can be focussed on a single task (Beggs and Plenz, 2003; Irrmischer et al., 2018b). A reduction in such psychological flexibility is a commonly reported depressive symptom and is a risk factor for serious depressive episodes (Coifman and Summers, 2019; Miranda et al., 2012; Stange et al., 2017). It is not clear, however, if these psychological symptoms can be mapped to underlying dynamic states. This would be an important step towards better understanding depressive symptoms at the level of large-scale neural systems (Deco et al., 2015). A symptom of particular interest from this point of view is negative rumination. This is a mental process involving repetitive thoughts about one’s feelings and problems (Smith and Alloy, 2009) that in depression is predominantly maladaptive. Excessive rumination can impact other thought processes and interfere with daily functioning (Lane and Northoff, 2017), being associated with deficits in problem-solving (Donaldson and Lam, 2004), attentional control (Lyubomirsky et al., 2003), and impacts upon memory (Liu et al., 2017).

Given the presence of fixed psychological states in depression, such as those seen in maladaptive rumination, we could hypothesise a similar fixedness in brain activity properties in people with high levels of depressive symptoms. This would be reflected in a reduced reactivity of the dynamic cortical state, meaning that it would not shift between critical and sub-critical states as normal when the participant changes their overall behavioural condition. Notably, the self-related thoughts that are a feature of rumination have been associated with the brain’s spontaneous activity (Huang et al., 2016; Nejad et al., 2013; Qin et al., 2016), which is altered in depression (Brakowski et al., 2017; Zhong et al., 2016) and which instantiates many features of cortical dynamics. This link further suggests a correspondence between fixed psychological states and reduced intrinsic activity reactivity.

A group of participants with a range of depressive symptom levels was therefore recruited to test whether differences in intrinsic activity reactivity were linked to depressive cognitive inflexibility. Following a dimensional approach (Insel et al., 2010), data from healthy participants were combined with data from people diagnosed with major depressive disorder (MDD). All participants had EEG recordings made during eyes-open and eyes-closed rest. DFA exponents were then calculated and compared between the two behavioural states.

This DFA change, taken as a proxy for the shift in criticality, was related to reported symptoms. A smaller difference in eyes-closed versus eyes-open DFA, taken to reflect a reduced flexibility in the cortical state, was expected to be correlated with greater reported depressive symptoms and with increased levels of negative rumination.

## Methods

### Participants

Thirty-seven healthy participants and twenty-six patients with MDD took part in the experiment. Three patients were excluded as they did not fully complete the psychological scales (MDD n = 23), leaving a total study group of 60 participants. Age and sex did not differ between these groups (Table 1A). Patients were recruited from the Department of Psychiatry at Shuang-Ho Hospital, New Taipei City. Controls were recruited from the local community. All patients were diagnosed according to DSM-5 criteria for current episode MDD (American Psychiatric Association, 2013). Patients predominantly had mild to moderate depression, as assessed with the Montgomery–Åsberg Depression Rating Scale (MADRS), with two having severe depression at the time of the experiment. The majority of patients were medicated at the time of the study (22/23) and had a mean time since the onset of their current episode of 5.8 months (± 6.1 SD). Patients with comorbid Axis I disorders were excluded, as diagnosed by a senior psychiatrist and confirmed by the Mini International Neuropsychiatric Interview (Sheehan et al., 1998). All participants were informed about the purpose of the study and procedures before giving written informed consent. This study was approved by the Joint Institutional Review Board, Taipei Medical University, Taipei City, Taiwan (N201603080).

**Table 1.**
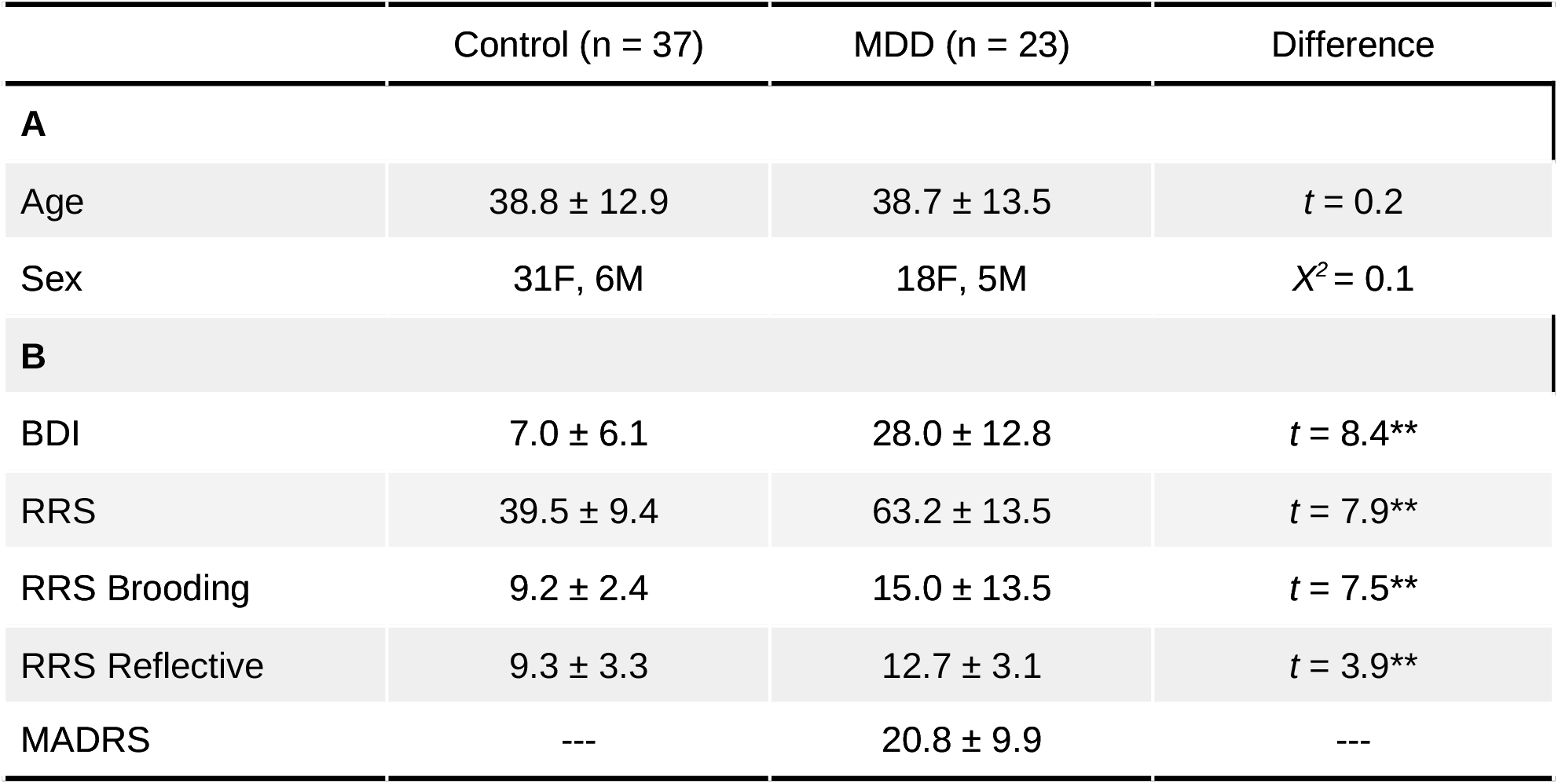
Demographics and questionnaires. Demographic details and questionnaire scores for the controls and MDD groups. Values given are group means ± standard deviation. Differences between the groups were compared with independent sample t-tests or chi-squared tests, as appropriate. ** denotes p < 0.001. BDI = Beck Depression Inventory, MADRS = Montgomery–Åsberg Depression Rating Scale, RRS = Rumination Response Scale.

### Psychological scales

All participants completed the Rumination Response Scale (RRS; Treynor et al., 2003) and the Beck Depression Inventory II (BDI; Beck et al., 1996). Both scales have been translated to Traditional Chinese and validated for use in Taiwan (Huang et al., 2015; Lu, 2002). Total scores for the RRS were calculated, along with Depression-related, Brooding, and Reflective rumination subscores (Treynor et al., 2003). The focus here was on the Brooding and Reflective subscores as these target ruminative behaviours that have less overlap with general depressive symptomatology (Treynor et al., 2003). Brooding rumination is generally taken to be maladaptive while Reflective rumination is a more positive behaviour (Smith and Alloy, 2009). MDD patients had higher scores than controls in all the scales measured (Table 1B).

### Participant grouping

All participants were combined into a single study group irrespective of MDD diagnosis. Associations with psychological scales were calculated across this full group. A median split was used to create “High BDI” and “Low BDI” groups where comparisons of people with higher or lower reported symptoms were required. The median BDI score was nine, with 31 participants having scores lower than or equal to this and 29 higher.

### EEG acquisition and preprocessing

EEG was recorded with Ag/AgCl electrodes mounted in an elastic cap (Easycap, Brain Products GmbH) using a 28-electrode arrangement following the International 10-20 System, including mono-polar electrodes (FP1/2, F7/8, F3/Z/4, FC3/Z/4/5/6, C3/Z/4, T7/8, CP1/Z/2/5/6, P3/Z/4/7/8, O1/Z/2) and four channels to record vertical and horizontal eye movements. Electrodes to record vertical movements were placed on the supraorbital and infraorbital ridges of the left eye, while those to record horizontal movements were placed on the outer canthi of the right and left eyes. A1 and A2 mastoid electrodes were used as an online reference (averaged across both). Online EEG was recorded with a BrainAmp amplifier (Brain Products GmbH). The data were sampled at 1000 Hz with a bandwidth of DC to 1000 Hz. Impedence was kept below 10kΩ at all electrodes. Data were recorded with BrainVision Recorder 1.2 software.

Participants sat in front of a computer screen with their chins on a chin-rest. During the eye-open session participants fixated a black fixation cross on a gray background. The instructions given to participants were to keep their eyes on the fixation cross, relax, and try to not fall asleep. The procedure during the eyes-closed session was the same but participants were instructed to keep their eyes closed. Eyes-open and eyes-closed data were both recorded for three minutes. The order was counterbalanced across participants. EEG recordings were made at approximately the same time of day for all participants (10-11am).

Data were preprocessed using EEGLAB (sccn.ucsd.edu/eeglab/index.php) and custom scripts running on MATLAB (MathWorks, version 2016b). The first thirty seconds of each acquisition were discarded to avoid activity related to the shifts between EO and EC states. The remaining EEG time series (150s) were FIR-filtered (1-45Hz band-pass, Blackman window with 1 Hz transition band) and then ICA was used to exclude eye movement induced artifacts. All signals were visually inspected to exclude any transient artifacts caused by head movements which were manually excluded and omitted from the subsequent analysis. All signals were then re-referenced to the common average.

### Detrended fluctuation analysis

DFA analysis was performed with the Neurophysiological Biomarker Toolbox (NBT, nbtwiki.net; Hardstone et al., 2012) toolbox for MATLAB. Preprocessed data were firstly band-pass filtered (FIR-filter) into the alpha (8-13 Hz) and beta bands (13-30 Hz). A Hilbert transform was then used to extract the amplitude envelopes in these bands for each electrode. Root-mean-square fluctuations of the integrated and linearly detrended envelope time series were calculated as a function of window size. Time windows corresponded to equally spaced points on a logarithmic scale of data lengths corresponding to 2-14 s for the alpha band and 1-14 s for beta (50% window overlap). These window sizes were selected to exclude temporal autocorrelations introduced by the band-pass filters (Irrmischer et al., 2018b). The DFA exponent is the slope of the line fitted through the fluctuation amplitudes against window lengths plotted on a logarithmic scale (Hardstone et al., 2012). Goodness-of-fit for these linear fits were calculated (R^2^) and compared between the EC and EO conditions. No differences were found at any electrodes.

### Statistical analysis

DFA values were first compared between the EC and EO conditions at each electrode. This was done for the alpha and beta bands using robust permutation tests of the difference in medians (5000 samples). Outlying values were excluded with reference to the median absolute deviation to the median, with a cut-off of 2.24 (Wilcox and Rousselet, 2018). P-values were adjusted for multiple comparisons across electrodes with Benjamini-Hochberg false discovery rate (FDR) correction (Benjamini and Hochberg, 1995), implemented in the statsmodels python toolbox (statsmodels.org). A threshold of *q* < 0.05 was applied.

EC-EO DFA differences were then related to BDI scores using robust regression to reduce the impact of outlying values on results (implemented in statsmodels). FDR correction was applied across all electrodes (*q* < 0.05). Electrodes where there was a significant association between EC-EO DFA differences and BDI scores were identified and mean EC-EO, EC, and EO DFA values across them calculated.

Using these mean values, EC-EO DFA differences were compared between the High and Low BDI score groups. This analysis aims to compare how reactive intrinsic activity properties were to the change in behavioural state. EC and EO DFA values were then compared separately between these same groups. Comparisons were done through robust permutation tests (5000 samples). Group medians are reported, along with their difference and its 95% confidence interval.

Finally, the relationship between beta DFA values at the same set of BDI-associated electrodes and rumination was investigated. EC-EO differences were firstly related to the Brooding and Reflective subscales of the RRS through a standardised multivariate robust regression. Separate regressions were then conducted to relate these subscales to EC and EO DFA values. The purpose of these regression analyses was to establish if DFA values correspond with one of these rumination styles specifically. Standardised model beta coefficients and their bootstrapped 95% confidence intervals are reported, along with coefficient *t*-statistics and *p*-values.

## Results

### Behavioural state change effect on DFA

Alpha and beta band DFA values were compared between the EC and EO conditions at each electrode. As shown in Figure 1, the switch to EO produced significant DFA reductions in both frequency bands. Alpha band changes were primarily found in frontal and occipital areas, extending through the midline (Figure 1A). Reductions in beta DFA were seen across all electrodes, with the largest at frontal and posterior electrodes and the smallest at electrodes associated with sensorimotor cortex (Figure 1B). DFA values at the electrodes with significant changes are illustrated in Figures 1C&D. These show that all DFA exponents lie between 0.5 and 1, indicating that LRTCs exist within the data (Hardstone et al., 2012).

**Figure 1:**
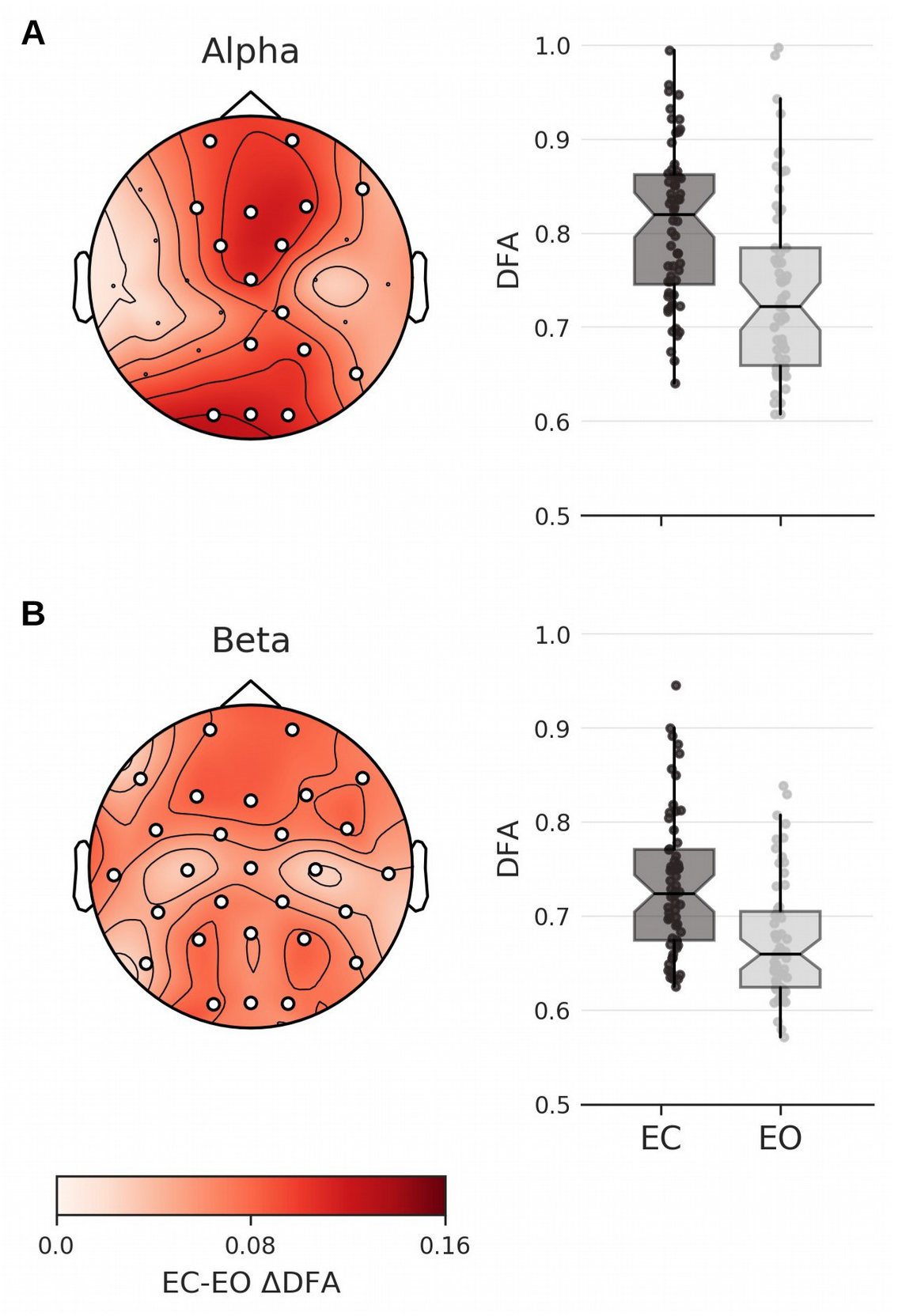
Topographical maps of EC-EO DFA difference. (A) EC alpha band DFA values were higher than EO across frontal and occipital electrodes, extending through the midline. Corresponding box plots illustrate this difference, showing average DFA values per participant across these electrodes. (B) EC beta band DFA values were higher than EO at all electrodes. This difference is illustrated in the corresponding boxplots. Significant electrodes after correction for multiple comparisons are indicated in white (*p*_FDR_ < 0.05). Topographic plots were made with the MNE toolbox (Gramfort et al., 2013).

### DFA changes and depressive symptoms

The change in DFA between EC and EO was then related to reported depressive symptoms, as measured with the BDI, through robust regression at all electrodes. No significant relationship between BDI scores and DFA differences was found at any alpha band electrode (Figure 2A). Beta DFA changes had a negative relationship with BDI scores at posterior and midline electrodes (Figures 2B). This relationship was not moderated by MDD diagnostic state (*t* = 1.07, *p* = 0.28).

**Figure 2:**
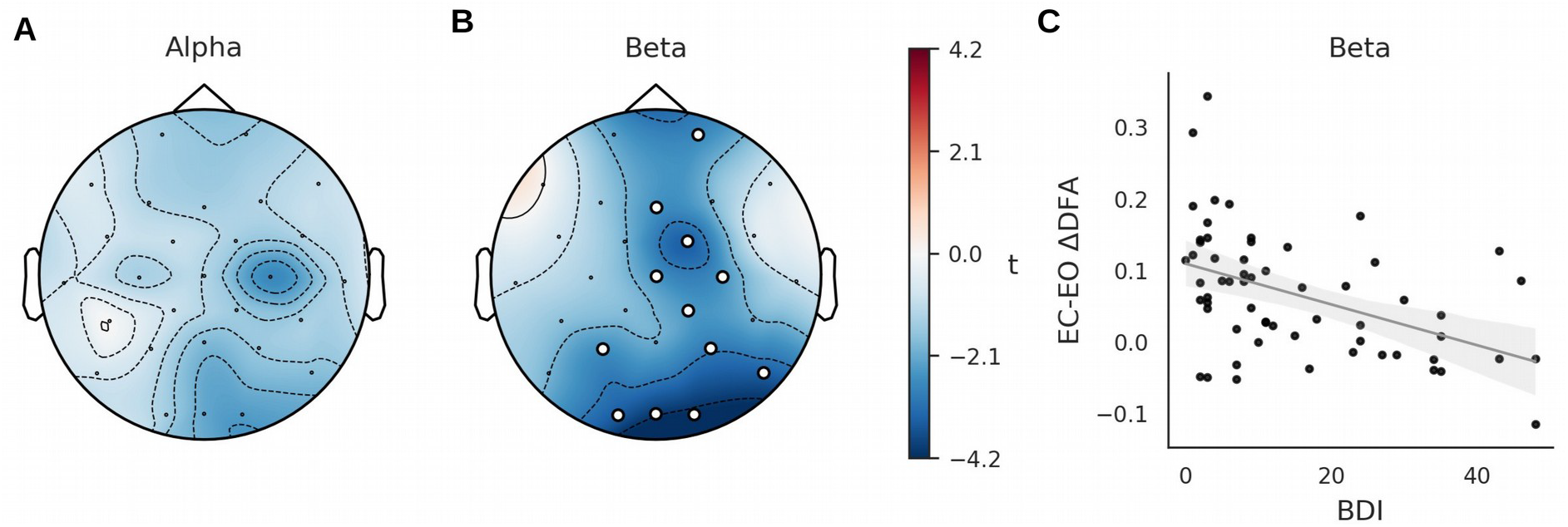
BDI and DFA value correlation. EC-EO DFA differences were regressed with BDI scores at each electrode. (A) No relationship was found with alpha band DFA differences. (B) A significant negative relationship was observed with beta DFA differences at occipital and midline electrodes. This is illustrated in a corresponding scatter plot (C) where mean DFA difference values across these significant electrodes are displayed against BDI scores. Significant electrodes after correction for multiple comparisons are indicated in white (*p*_FDR_ < 0.05). BDI = Beck Depression Inventory, DFA = detrended fluctuation analysis, EC = eyes closed, EO = eyes open. Beta DFA values were averaged across the electrodes associated with BDI scores and then compared between the Low (n = 31) and High BDI groups (n = 29). Participants with low BDI scores showed a larger EC-EO difference than those with higher ones (Low *M* = 0.09, High *M* = 0.02, Δ*M* = 0.071, 95% CI = [0.048 0.13], *p*_FDR_ < 0.001, Figure 3A). DFA changes for the Low BDI group differed from zero (*p*_FDR_ < 0.001) while those for the High group did not (*p*_FDR_= 0.093).

As shown in Figure 3B, DFA values in the EC condition were greater for the Low than the High BDI group (Low *M* = 0.76, High *M* = 0.69, Δ*M* = 0.076, 95% CI = [0.031 0.14], *p*_FDR_ = 0.004). The same effect was not seen in the EO condition, where DFA values did not differ between groups (Low *M* = 0.67, High *M* = 0.67, Δ*M* = −0.006, 95% CI = [−0.037 0.031], *p*_FDR_ = 0.48).

**Figure 3:**
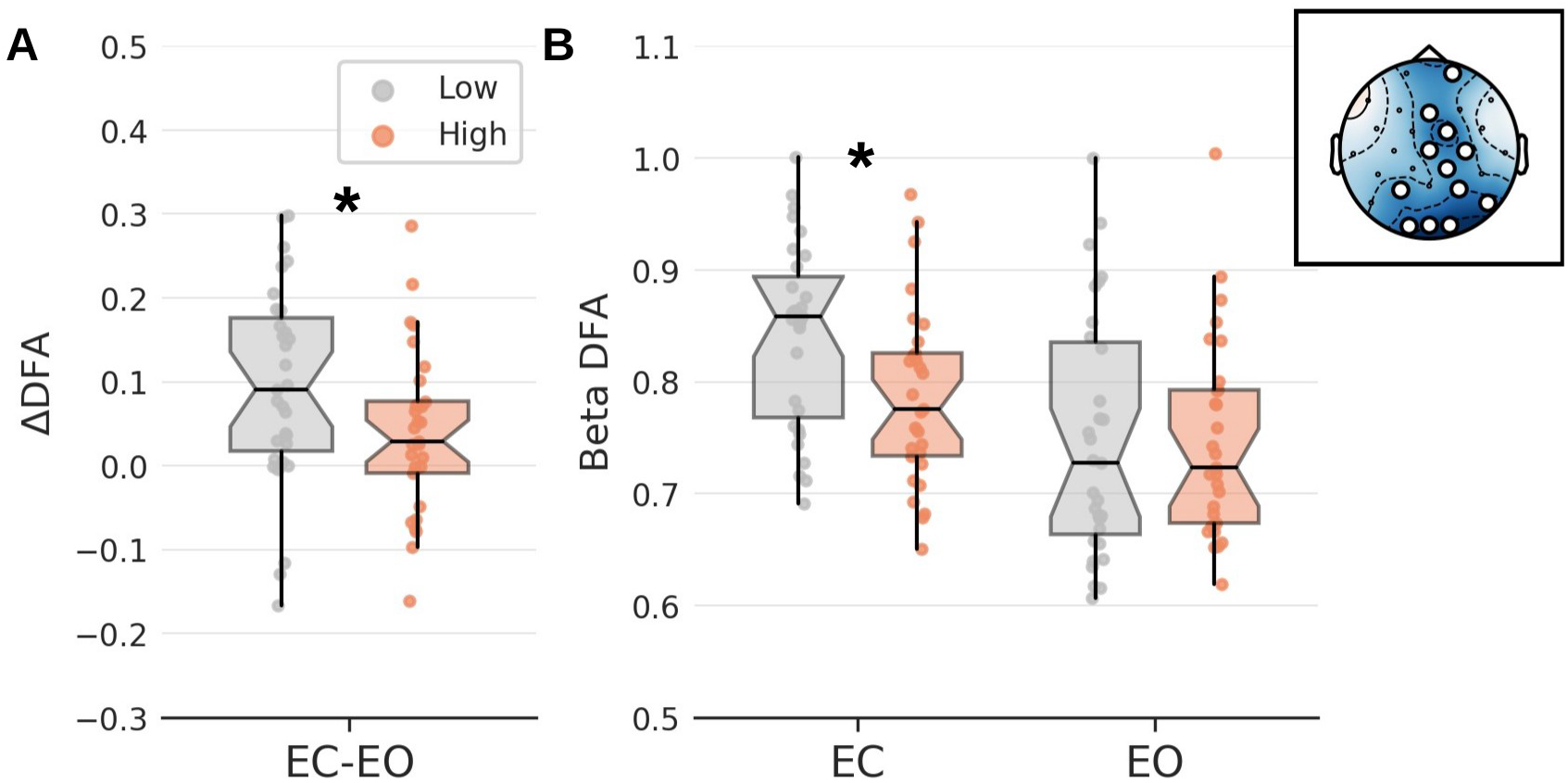
BDI and DFA value correlation. EC-EO DFA differences were regressed with BDI scores at each electrode. (A) No relationship was found with alpha band DFA differences. (B) A significant negative relationship was observed with beta DFA differences at occipital and midline electrodes. This is illustrated in a corresponding scatter plot (C) where mean DFA difference values across these significant electrodes are displayed against BDI scores. Significant electrodes after correction for multiple comparisons are indicated in white (p_FDR_ < 0.05). BDI = Beck Depression Inventory, DFA = detrended fluctuation analysis, EC = eyes closed, EO = eyes open.

### Beta DFA and rumination

The relationship between DFA values at electrodes associated with BDI scores and RRS subscales was investigated through multivariate robust regression. EC-EC changes were not associated with rumination scores (Brooding - β = −0.3, 95% CI = [−0.64 0.07], *t* = −1.67, *p*_FDR_= 0.29; Reflective - β = 0.03, 95% CI = [−0.3 0.41], *t* = 0.16, *p*_FDR_ = 0.87). A negative association with EC DFA values was, however, observed for Brooding rumination (β = −0.53, 95% CI = [−0.86 −0.15], *t* = −2.86, *p*_FDR_ = 0.012; Figure 4A). This was not seen with Reflective rumination scores (β = 0.22, 95% CI = [−0.15 0.56], *t* = 1.18, *p*_FDR_ = 0.36; Figure 4B). No such relationship was found between EO DFA values and either Brooding (β = −0.21, 95% CI = [−0.67 0.19], *t* = −1.12, *p*_FDR_ = 0.69; Figure 4C) or Reflective rumination (β = 0.08, 95% CI = [−0.25 0.44], *t* = 0.4, *p*_FDR_ = 0.69; Figure 4C). The relationship between EC DFA and Brooding rumination was not moderated by Low or High BDI group membership (*t* = 1.7, *p* = 0.09).

**Figure 4:**
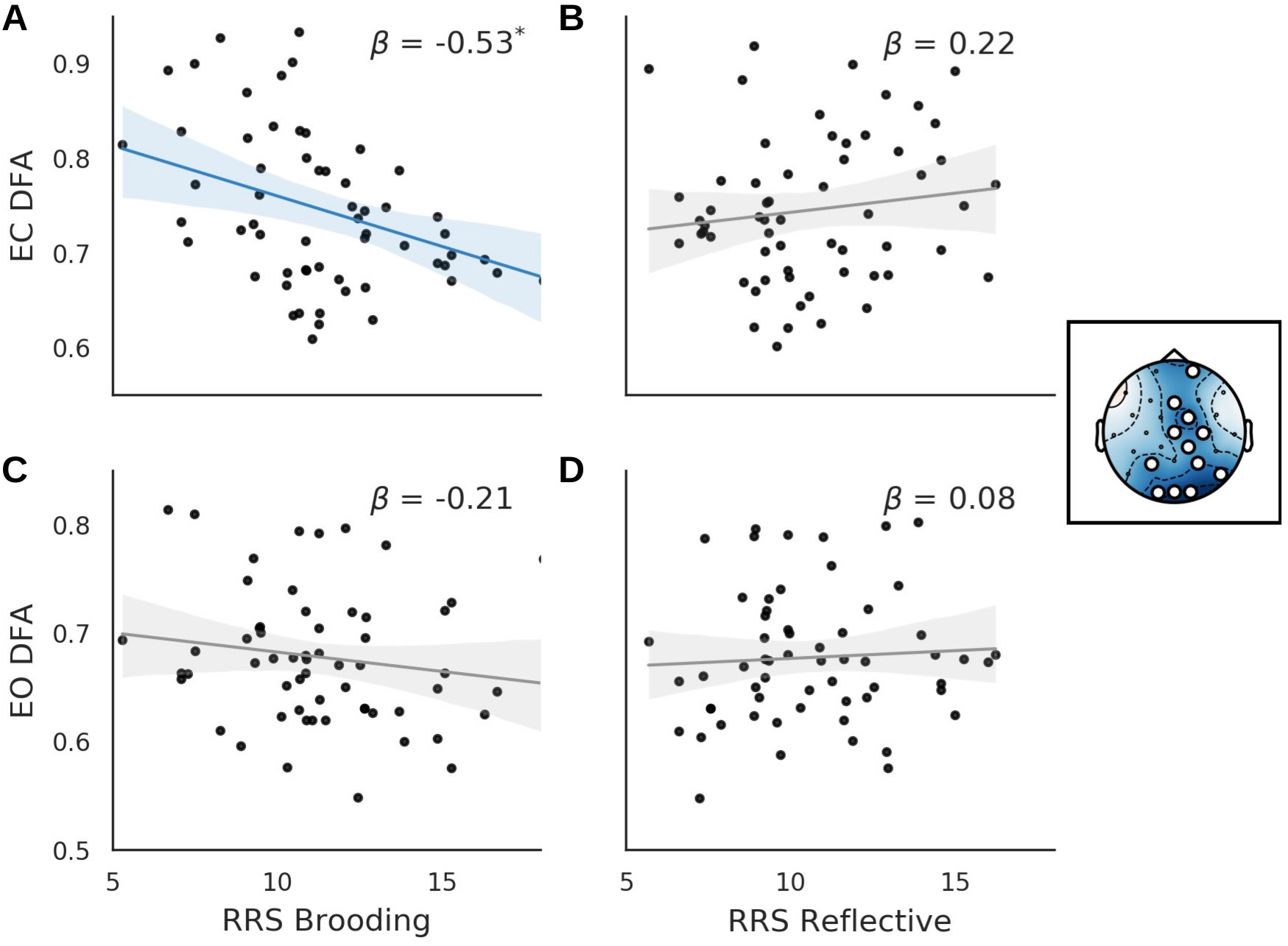
DFA values and rumination. (A) A negative correlation between beta band EC DFA values and the Brooding subscale of the RRS was seen. No such relationship was found with the Reflective subscale (B). EO DFA values were not related to either subscale (C & D). Note that scatter plot values are adjusted for the corresponding RRS subscores and so the position of specific data points on the different axes may differ. The electrodes from which DFA values were averaged are shown in the topographic inset. * denotes *p*_FDR_< 0.05. DFA = detrended fluctuation analysis, EC = eyes closed, EO = eyes open, RRS = Rumination Response Scale.

## Discussion

Detrended fluctuation analysis was used to quantify long-range temporal correlations within EEG activity recorded at rest with the eyes open and closed. These DFA values were found to reduce in the EO condition when compared to EC in both the alpha and beta frequency bands. The change in DFA seen in the beta band correlated with reported depressive symptoms at occipital and midline electrodes. Spliting participants into high and low depressive symptom groups, it was found that DFA values at these electrodes did not change between EC and EO for the latter group. This was due to EC DFA values remaining at the same level as in the EO condition for these participants. Finally, it was shown that these beta band EC DFA values specifically correlate with brooding rumination.

The reduction in DFA seen in the EO condition when taking the participants as a whole mirrors the LRTC reductions that occur when people engage in a particular task, as compared to a task-free state (He, 2014). This effect has been reported for a number of imaging modalities, including EEG (He, 2011; Irrmischer et al., 2018b, 2018a), and across frequency ranges (He, 2011; He et al., 2010). A reduction in LRTC reflects a system operating closer to sub-critical levels, meaning that activity has less temporal complexity and can dwell more consistently close to a particular configuration (Irrmischer et al., 2018b). This contrasts with the greater degree of LRTC indicated by the higher DFA values seen in the EC state. Such increased LRTC are likely to reflect brain activity operating near criticality (Deco and Jirsa, 2012), where information handling and the capacity to respond to varied inputs are optimised (Gautam et al., 2015; Hahn et al., 2017; Shew et al., 2011).

The change from EC to EO has been shown previously to produce extensive alterations to brain activity patterns (Barry et al., 2007; Marx et al., 2003). This occurs in a manner that is partly independent of the change in visual stimulation (Marx et al., 2003), indicating that the behavioural change itself produces a shift in activity structure. These changes can be seen within a wider context of brain activity reorganisations brought about by changes in behavioural context (Clancy et al., 2019; Marques et al., 2020) and the manner in which these can optimise context-dependant functions (Benjamin et al., 2018; Cao and Händel, 2019). The observed negative correlation between BDI scores and beta band EC-EO DFA changes may therefore reflect a reduced capacity to appropriately shift brain state based on behavioural context with increasing depressive symptoms.

When split into Low and High BDI groups, people with higher depressive symptoms did not show increases in beta DFA values in the EC condition. An association between depressive symptoms and EC beta activity has been noted in a number of previous studies (Newson and Thiagarajan, 2019), including ones specifically investigating LRTC (Lee et al., 2007; Linkenkaer-Hansen et al., 2005). A reduction in beta LRTC is also seen in other brain disorders, including Alzheimer’s disease (Montez et al., 2009) and schizophrenia (Nikulin et al., 2012), although with different spatial distributions. Altered activity dynamics are also seen in Parkinson’s disease (Hohlefeld et al., 2012), epilepsy (Monto et al., 2007), and autism (Lai et al., 2010). This suggests that a disruption to the temporal structure of brain activity may be a common feature of pathological conditions (Hardstone et al., 2012), with symptomatic variation arising from the specific brain networks and frequency ranges within which such changes occur (Northoff and Duncan, 2016).

LRTC levels remaining lower in the EC state for the High BDI group means that those with higher depressive symptoms show less of a transition from a sub-critical state towards a critical one. This implies that their brain will display lower neural state dynamics over time which may in turn lead to less flexibility in mental processing. Notably, inflexibility across a number of psychological domains is a common feature in those with depression and has been identified as a risk factor for developing such a condition (Coifman and Summers, 2019; Miranda et al., 2012; Stange et al., 2017). This highlights the relevance of investigating such functions and related brain features across a full range of depressive symptom levels in a dimensional fashion.

Of particular interest in the context of the current work are attentional inflexibilities (Fergus et al., 2013) as beta band activity, such as that highlighted here, has been linked to the executive and cognitive control processes associated with them (Schmidt et al., 2019; Stoll et al., 2016). This EC beta band activity was found here to correlate with brooding rumination, with no such correlation found with the more beneficial reflective rumination. Negative rumination of this sort is of central relevance to depression as it is associated with worse outcomes (Huffziger et al., 2009; Spinhoven et al., 2018; Surrence et al., 2009) and represents a risk factor for developing depression in healthy individuals (Spasojević and Alloy, 2001). Interestingly, no correlation was found with EO DFA values, nor with EC-EO differences, suggesting that the association with EC beta DFA may not result from a general effect of more rumination leading to lower DFA values across all behavioural states. Instead we may be observing a situation whereby the neural inflexibility indicated by the lack of EO to EC DFA change predisposes people to engage in negative rumination by retaining an overly attentive state. Where visual or other external input is reduced, this attentive state may instead orient to self-oriented content, such as is the focus of rumination (Nejad et al., 2013). This supposition is speculative but does present a target at the level of large-scale neural systems for further investigations of specific depressive symptoms. In addition, the suggested relationships also point to potential clinical uses for behavioural interventions such as mindfulness training and exercise that improve the flexibility of brain activity (Gärtner et al., 2017; Irrmischer et al., 2018a; Wu et al., 2019).

A number of limitations of this work must be noted. Firstly, although the sample size used was relatively large (n = 60), it will be important to replicate the results in an independent sample in the future. Secondly, the majority of the MDD patients who took part were medicated with drugs targeting a variety of neurotransmitter systems. Given the variability in systems targeted, it is unlikely that the relationship found between behaviour and EEG properties across the full patient plus control dataset are driven by medication effects but this cannot be ruled out and so studies on unmedicated patients would be justified. This would be particularly important when testing particular models of neurotransmitter changes that may be driving ruminative behaviours. Thirdly, the majority of the participants in the study were female and so it is possible that the findings are sex-specific. Unfortunately, there are insufficient participants here to test this and so future research is required to confirm the generalisability of the results. As well as this, the menstrual cycle has been suggested to influence different aspects of intrinsic brain activity and so this factor may impact the reported results (Duncan and Northoff, 2012; Syan et al., 2017; Weis et al., 2019). Relevant details from participants were not available to test any association between cycle stage and rumination or brain activity and so this remains an important outstanding research question.

To conclude, DFA was used to quantify LRTC in neural activity during rest with the eyes open and closed in a group of healthy participants and patients with MDD. Depressive symptoms were related to an inability to shift between a critical and sub-critical state between the EC and EO state, with those with more symptoms remaining in a state with lower DFA values. We can speculate that this may reflect a general cognitive inflexibility resulting from a lack in neural activity reactivity that may predispose people to overly engage in self-directed attention such as is seen in negative rumination.

## Acknowledgements

The authors would like to thank all participants for their time and effort. They are grateful also to Ching Lin for help with patient recruitment and data collection. This work was supported by funding from the Taiwan Ministry of Science and Technology to TJL (104-2420-H-038-001-MY3; 105-2632-H-038-001-MY3), TYH (106-2410-H-038-004-MY2; 104-2410-H-038-012-MY2), and NWD (108-2410-H-038-008-MY2).

## Competing interests

The authors declare no conflicts of interest.

## Author contributions

NWD and TYH analysed the data; PZC, HYW, and HCL collected the data; NWD, TYH and TJL designed the study; NWD and TYH wrote the manuscript; all authors revised and approved the manuscript.

## Data accessibility

The data included in this analysis are available at https://osf.io/e7mwz/.

## Abbreviations

BDI: Beck Depression Inventory II
EEG: electroencephalogram
EC: eyes-closed
EO: eyes-open
DFA: detrended fluctuation analysis
FDR: false discovery rate
LRTC: long-range temporal correlations
MDD: major depressive disorder
MADRS: Montgomery–Åsberg Depression Rating Scale
RRS: Rumination Response Scale

## Notes

#### Summary of Updates

Restructured to improve clarity.

